# Generating immunogenomic data-guided virtual patients using a QSP model to predict response of advanced NSCLC to PD-L1 inhibition

**DOI:** 10.1101/2023.04.25.538191

**Authors:** Hanwen Wang, Theinmozhi Arulraj, Holly Kimko, Aleksander S. Popel

## Abstract

Generating realistic virtual patients from a limited amount of patient data is one of the major challenges for quantitative systems pharmacology modeling in immuno-oncology. Quantitative systems pharmacology (QSP) is a mathematical modeling methodology that integrates mechanistic knowledge of biological systems to investigate dynamics in a whole system during disease progression and drug treatment. In the present analysis, we parameterized our previously published QSP model of the cancer-immunity cycle to non-small cell lung cancer (NSCLC) and generated a virtual patient cohort to predict clinical response to PD-L1 inhibition in NSCLC. The virtual patient generation was guided by immunogenomic data from iAtlas portal and population pharmacokinetic data of durvalumab, a PD-L1 inhibitor. With virtual patients generated following the immunogenomic data distribution, our model predicted a response rate of 18.6% (95% bootstrap confidence interval: 13.3-24.2%) and identified CD8/Treg ratio as a potential predictive biomarker in addition to PD-L1 expression and tumor mutational burden. We demonstrated that omics data served as a reliable resource for virtual patient generation techniques in immuno-oncology using QSP models.

## INTRODUCTION

Lung cancer is the top leading cause of cancer death in the U.S. with 130,180 estimated deaths in 2022 *(1)*. Non-small cell lung cancer (NSCLC) is the most common subtype of lung cancer, which accounts for about 84% of total lung cancer cases *(2)*. Since 2015, immune checkpoint inhibitors targeting programmed cell death protein (death-ligand) 1 [PD-(L)1] and cytotoxic T-lymphocyte antigen-4 (CTLA-4) have begun to receive approval from the U.S. Food and Drug Administration (FDA) for advanced NSCLC. For patients without actionable mutations [i.e., epidermal growth factor receptor (EGFR) and anaplastic lymphoma kinase (ALK)], different immune checkpoint inhibitors are recommended in single-agent or dual immunotherapy, or in combination with chemotherapy or bevacizumab, an anti-VEGF antibody, based on PD-L1 expression level on tumor cells *(3)*. PD-L1 expression, as a regulator of antitumor response, an indicator of T cell infiltration into the tumor, and the target of immune checkpoint inhibitors, has been widely used as a predictive biomarker for immunotherapy in advanced NSCLC *(4)*. Although immunotherapy has significantly improved the overall survival rate in advanced NSCLC when compared to conventional treatments, less than half of the patients respond (including those with >50% PD-L1 expression on tumor cells), and the 3-year survival rate is significantly lower in patients without actionable mutations *(5)*. Due to the low prevalence of actionable mutations *(6)*, novel combination regimens that involve immune checkpoint inhibitors are under investigation in clinical trials *(3)*.

In conjunction with the clinical effort, quantitative systems pharmacology (QSP) models have been developed in the past few years, aiming to predict clinical benefits of treatment of interest in complex diseases like NSCLC. QSP integrates mechanistic knowledge from multiple disciplines, such as systems biology, (patho)physiology, and pharmacology, and investigates dynamic behavior of a system as a whole *(7)*. Particularly in immuno-oncology, increasing number of QSP models was developed to study drug exposure-efficacy and exposure-toxicity relationships, predict efficacy, and identify predictive biomarkers for newly discovered drugs, including T cell engagers, immune checkpoint inhibitors (ICIs), and chimeric antigen receptor (CAR) T cells *(8)*. The main goal is to assist drug and clinical trial designs, such as target and dose optimization, and to reduce the cost and time in drug development *(9, 10)*. Among these efforts, our previously developed QSP platform, QSP-IO, has been applied to simulate tumor response to ICIs and their combinations with other types of treatment in early-stage NSCLC *(11, 12)*, breast cancer *(13–15)*, colorectal cancer *(16, 17)*, and hepatocellular carcinoma *(18)*. Although recent QSP models provide reliable efficacy predictions for clinical trials at the population level, one of the major challenges remains in virtual patient generation, which aims to generate virtual patient cohorts that represent the interindividual variabilities observed in real-world data while falling within the physiologically plausible ranges *(15)*.

Although methods have been proposed to guide virtual patient generation, they have not been widely applied to large-scale models like QSP *(19, 20)*. The focus of this study is therefore to investigate the performance of published virtual patient generation methods when integrated with our latest QSP-IO platform *(15)*. Specifically, we applied two virtual patient generation methods: 1) probability of inclusion *(21)* to select virtual patients that statistically match patient data from the Cancer Research Institute (CRI) iAtlas, a platform storing results from immunogenomic analyses of TCGA data *(22)*; and 2) compressed latent parameterization *(23)* to generate pharmacokinetic (PK) parameters based on pseudo-patient level data from population PK analysis of durvalumab, a PD-L1 inhibitor. For model validation, we first predicted the objective response rate of the generated virtual patients to durvalumab to compare with results from Study 1108 (NCT01693562), a phase 1/2 clinical trial in advanced NSCLC *(24)*. In addition, we validated model-predicted immune cell densities against results from a digital pathology analysis, which is a quantitative analysis of histological images that provides spatial densities of immune markers of interest, such as CD4, CD8, and FoxP3, in different tumor regions *(25–29)*.

## RESULTS

### Model parameterization

Figure 1 illustrates the workflow of the present analysis. We utilized our previously developed QSP platform *(15)* that describes the cancer-immunity cycle *(30)* and recalibrated the cancer-type specific parameters using experimental and clinical data on NSCLC. Table 1 lists the recalibrated parameters with the data we used to estimate their values. As we aim to simulate a phase 1/2 clinical trial of durvalumab that enrolled patients with stage III NSCLC, data on stage III NSCLC were preferentially used when available. Overall, the model involves four main compartments: central, peripheral, tumor, and tumor-draining lymph node. Ten modules were incorporated to investigate dynamics of cellular and molecular species, including cancer cells, T cells (i.e., effector, helper, and regulatory T cells), immune checkpoints, and durvalumab, in their corresponding compartments. Since majority of the clinical data for virtual patient generation and model validation (e.g., CD8 and CD4 T cell density) were collected from tumor samples, the model was best trained to describe immune cell dynamics in the tumor compartment, which is therefore the focus of the following analyses. With the recalibrated model, we first generated 30,000 plausible patients, 629 of which were selected to form a virtual patient cohort. The selection process was guided by immune cell subset ratios (i.e., M1/M2, CD8/Treg, and CD8/CD4) estimated from immunogenomic analysis, as described below in Methods.

**Figure 1.**
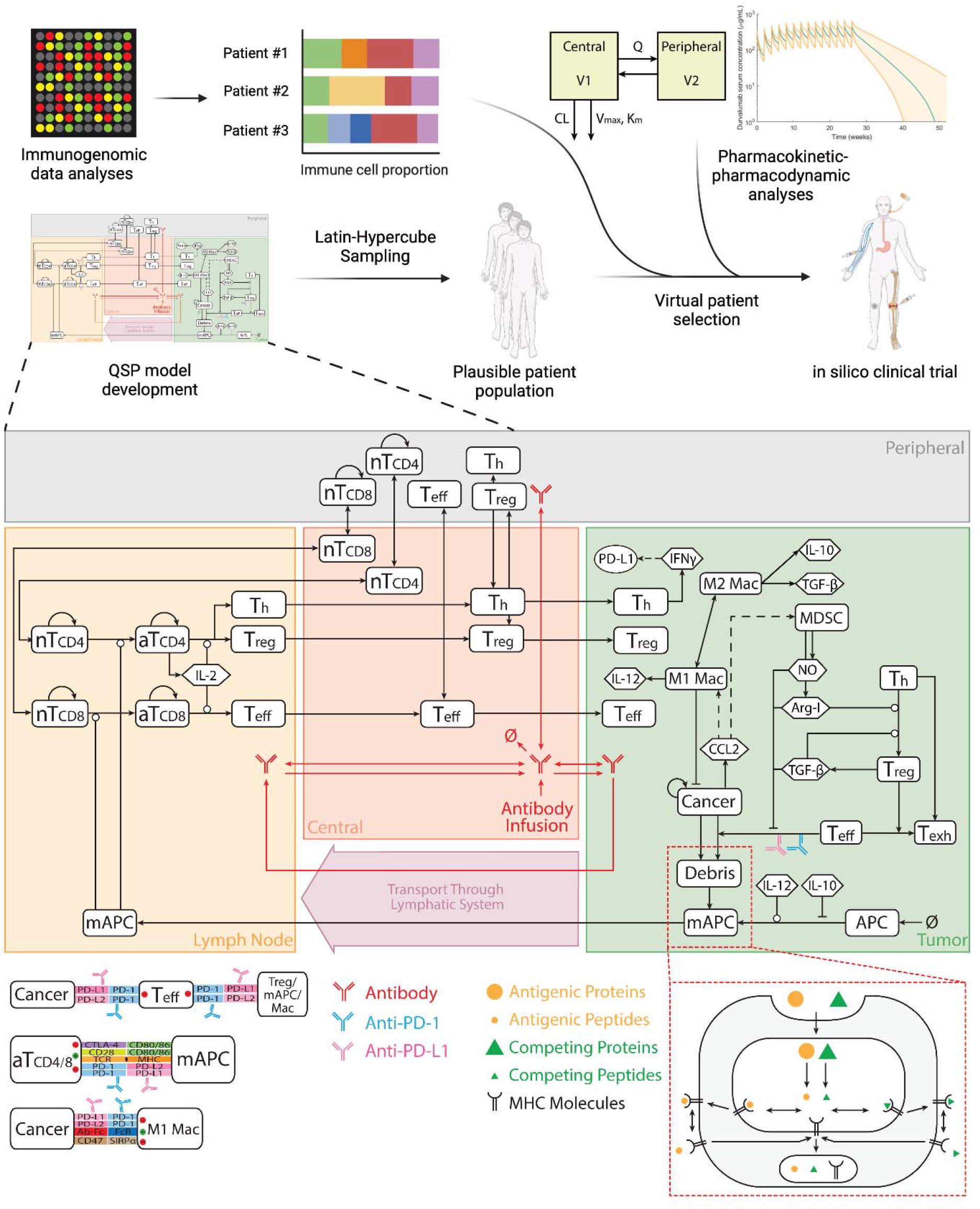
Paradigm of immunogenomic data-guided virtual patient generation and in silico clinical trial simulation using a mechanistic quantitative systems pharmacology model. The model is comprised of four compartments: central, peripheral, tumor, and tumor-draining lymph node, which together describe the cancer-immunity cycle. nT, naïve T cell; aT, activated T cell; NO, nitric oxide; Arg-I, arginase I; Treg, regulatory T cell; Teff, effector T cell; Th, helper T cell; Mac, macrophage; mAPC, mature antigen presenting cell. Cytokine degradation and cellular clearance are omitted in the figure.

**Table 1.**
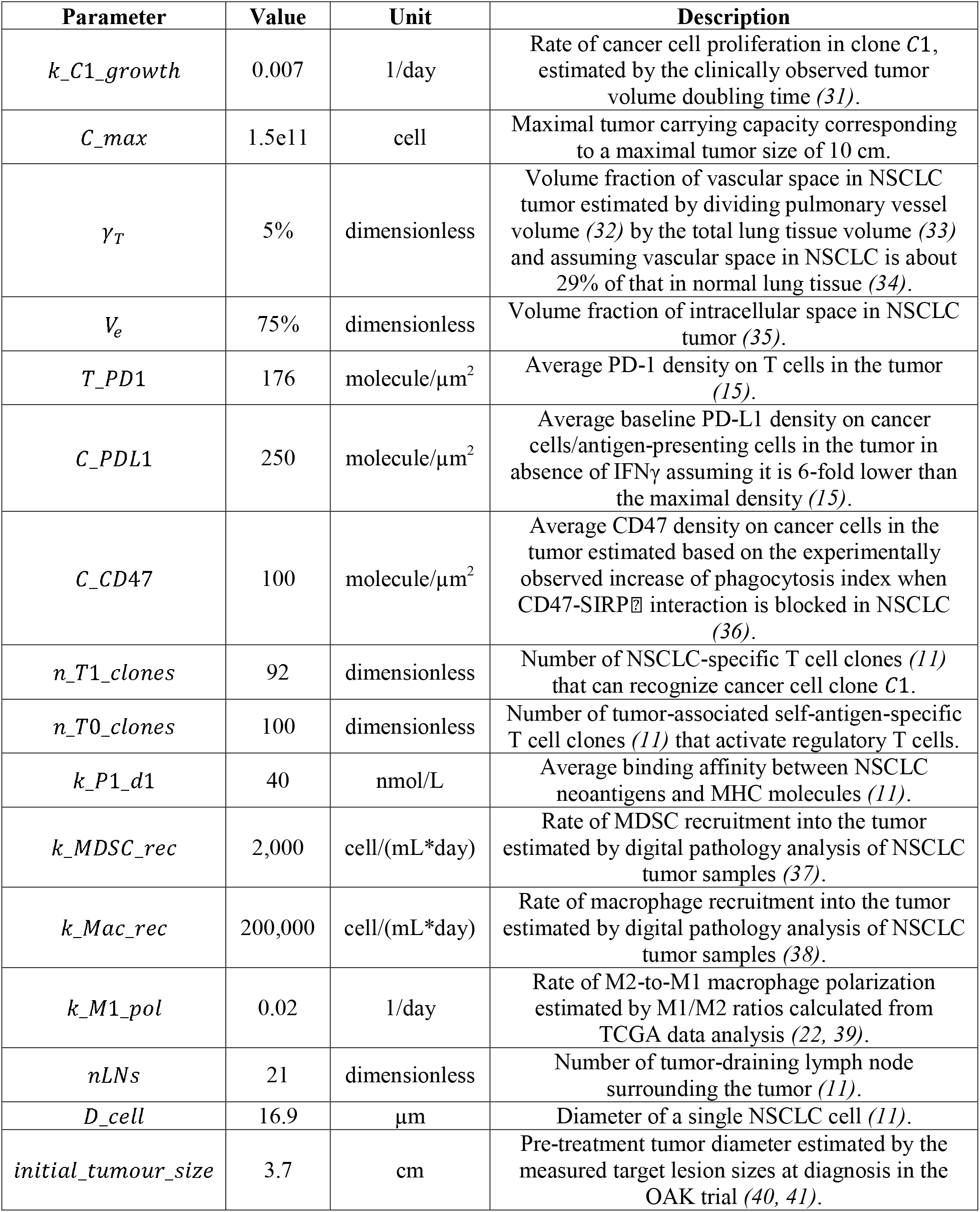
Non-small cell lung cancer (NSCLC)-specific parameters in the quantitative systems pharmacology model. MHC, major histocompatibility complex.

### Virtual patient generation

We first confirmed that the virtual patient cohort statistically matched the immunogenomic data that guided the selection process. Figure 2A shows the estimated probability densities of the three immune subset ratios in: 1) the plausible patients, 2) the final virtual patient cohort, and 3) the immunogenomic dataset from iAtlas portal *(22)*. We also compared the distributions of the three ratios between the observed data and the virtual patients in Figure 2B. When comparing the distributions using Kolmogorov-Smirnov tests, the test statistics were 0.07, 0.06, and 0.06 with p-values of 0.30, 0.51, and 0.44, indicating that the virtual patient’s distributions were not statistically different from those of the immunogenomic data. Also shown in Figure 2A, the ranges of immune subset ratios in the plausible patients were wider than that in the immunogenomic data, which were narrowed by the selection process to generate the virtual patient population that better fitted to the data. Here, we use immune subset ratios instead of the proportions of immune cells in leukocytes as reported in iAtlas database because the proportions do not directly correspond to our model outputs, where immune cell densities are calculated in cells per cubic milliliter of tumor. In addition, we only selected data on M1/M2 macrophages, CD8, CD4, and regulatory T cells because other cellular types on the database (e.g., natural killer cells, B cells) were not explicitly represented in the current model.

**Figure 2.**
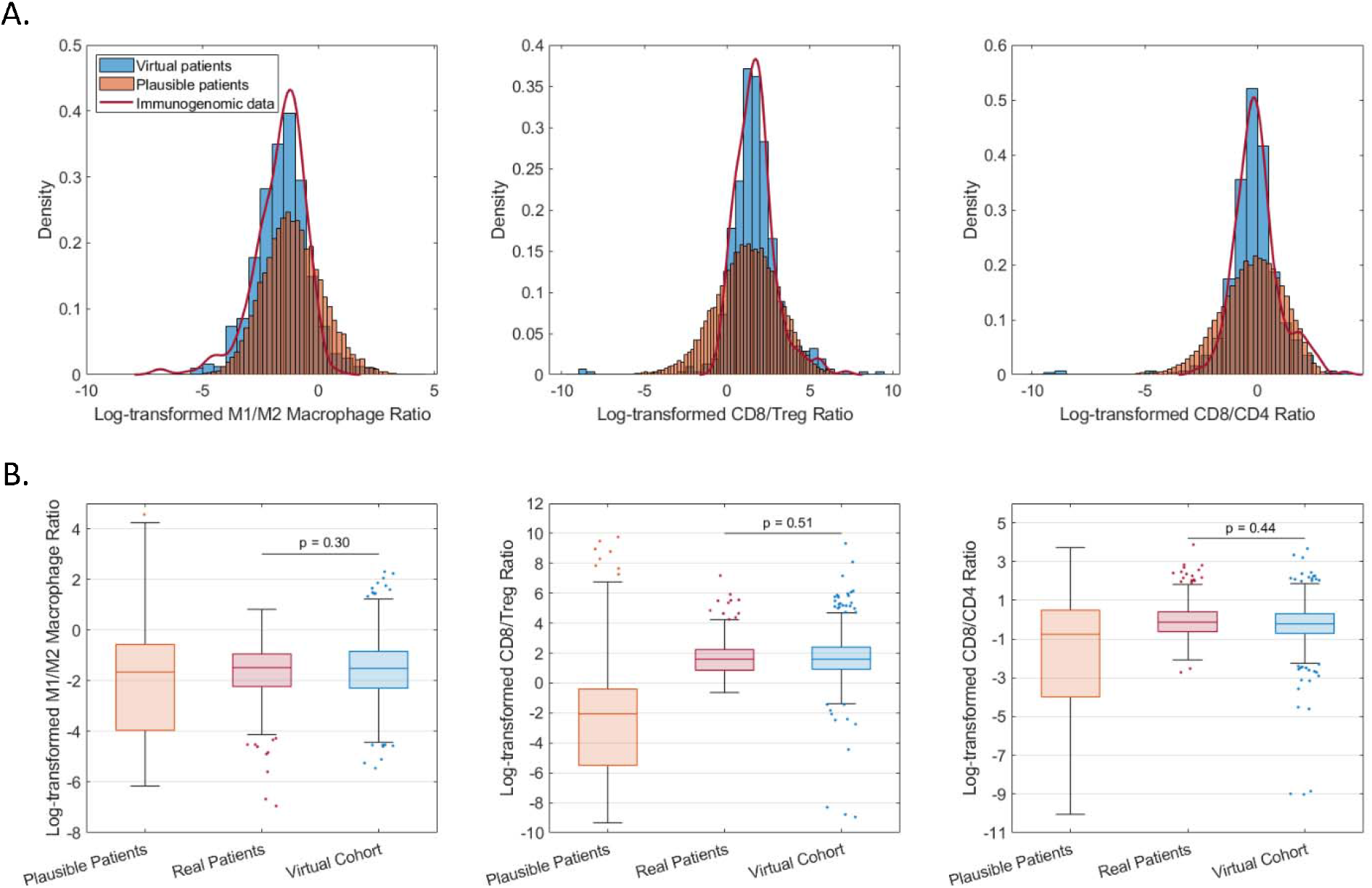
Probability density (A) and distribution (B) of model-predicted pre-treatment immune cell subset ratios in the virtual patient cohort compared with that calculated from immunogenomic data analysis. In the top panels, red lines represent the estimated probability density functions from immunogenomic data analysis; orange bars represent probability densities in the randomly generated plausible patient population; blue bars represent probability densities in the selected virtual patient cohort. In the bottom panels, 25^th^, 50^th^, and 75^th^ percentiles are encoded by box plots with whiskers that define 1.5 times the interquartile range away from the bottom or top of the box. Natural-log transformation was performed for the immune cell subset ratios during virtual patient generation.

Next, we validated the virtual patient cohort by comparing other pre-treatment characteristics of the virtual patients with observed data from clinical analyses. Figure 3 shows the probability densities of the pre-treatment tumor size, tumor doubling time, densities of CD8, CD4, Treg, and tumor-associated macrophages (TAMs), myeloid-derived suppressor cells (MDSCs), and PD-L1 expression in the tumor. The median tumor size is 3.7 cm with a range between 1.5 and 9.9 cm, which is consistent with the measurement from the OAK trial for stage IIIB/IV NSCLC *(40, 41)*. The tumor volume doubling time (TVDT) of virtual patients was calculated by *TVDT* = *δt* · log 2/(log *V*_*t*2_ − log *V*_*t*1_). *δt* is the time interval between two CT scans that are usually performed at diagnosis and at the beginning of the treatment. *V*_*t*1_ and *V*_*t*2_ are the measured tumor volumes at the two time points. Assuming a *δt* of 8 weeks, we estimated the mean TVDT to be 113 days with a median of 89 days in the virtual patients, which aligned with clinically measured TVDT of stage III NSCLC *(31)*.

**Figure 3.**
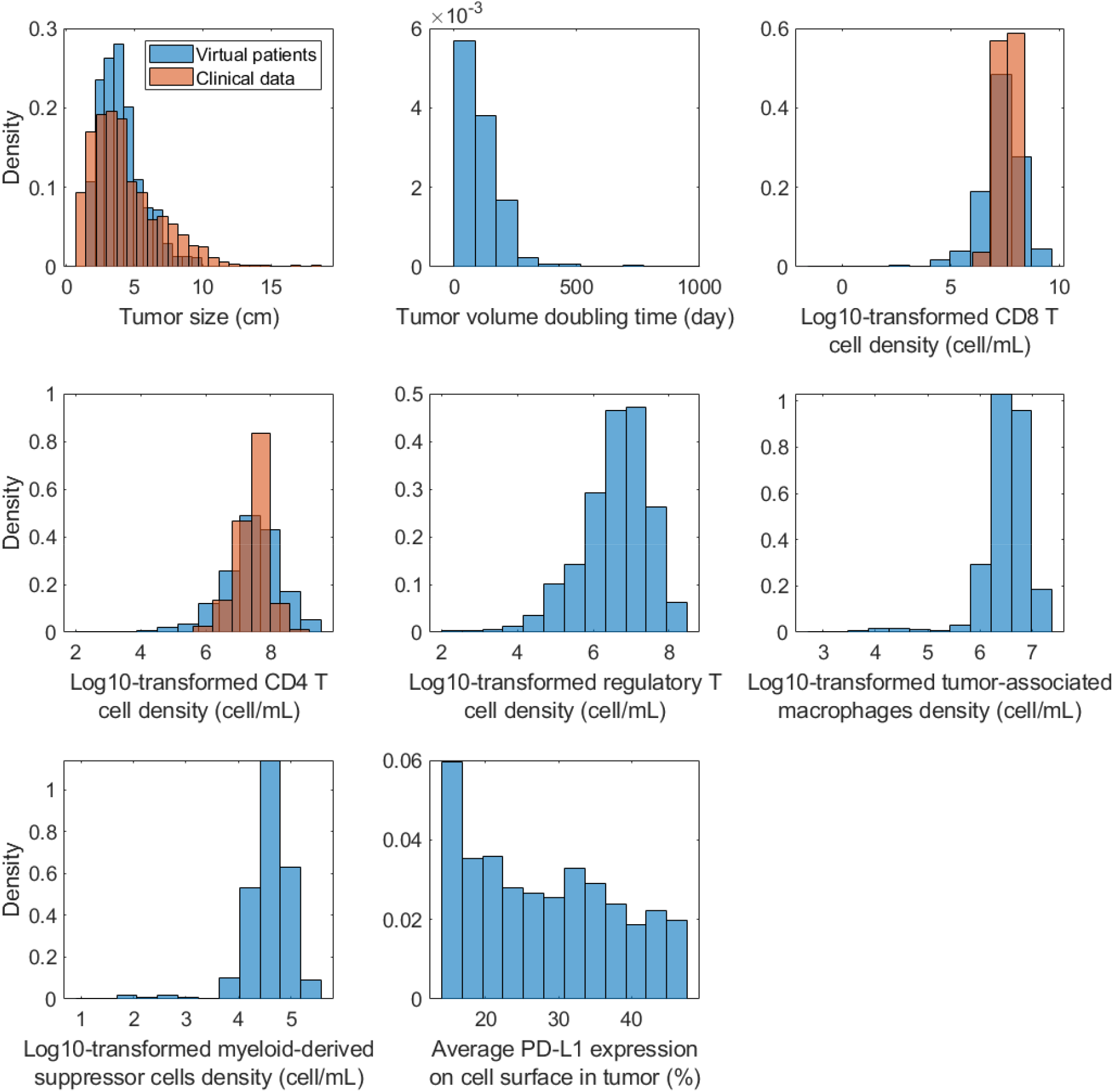
Probability density of model-predicted pre-treatment variable distribution in the virtual patient cohort. Clinical data on tumor size were from the OAK trial *(40, 41)*. CD8 and CD4 T cell densities in NSCLC tumor were obtained from Kilvaer et al. *(42)*. Average PD-L1 expressions on cell surface were estimated by dividing the model-predicted average PD-L1 density across all cells in the tumor by a theoretical maximal PD-L1 density of 1,770 molecules/μm^2^.

The median densities of CD8, CD4, Treg, TAMs, and MDSCs were 2.6e7, 3.2e7, 5.4e6, 3.7e6 and 4.2e4 cells/mL in the tumor, which were in agreement with multiple digital pathology analyses *(37, 38, 43–45)*. In Figure 3 and Supplementary Figure 1, we compared the distributions of CD8 and CD4 density between virtual patients and clinical data from patients with stage III NSCLC, which were obtained from Kilvaer et al. *(42)*. The conversion from a 2-D density from digital pathology analyses to the 3-D density was performed using equations presented in Mi et al. *(26)*. Notably, the ranges of CD8 and CD4 T cell densities in the virtual patients were wider than that in the clinical data, which was due to the inherent uncertainty resulted from model parameterization and virtual patient generation. Nonetheless, the model-predicted T cell subset densities likely fell within the physiologically reasonable ranges, as the proportions of CD8 and CD4 T cells ranged from 0% to 40% in iAtlas data (zeros were removed when calculating immune subset ratios to avoid singularities). Overall, the virtual patient cohort shows resemblance to real patient populations observed in clinical analyses of NSCLC.

### Variability in pharmacokinetic parameters

According to the population PK (popPK) study of durvalumab, drug exposure can be affected by characteristics like body weight, serum albumin, and soluble PD-L1 level *(46)*. Although most of these clinically measured characteristics are not present in the QSP model, the covariate effect on PK was included during reproduction of the PK data to be fitted with the QSP model. Since immunogenomic data used to select virtual patients above were not coupled with PK-related data, we independently generated PK parameters for the virtual patients via compressed latent parametrization *(23)*. This optimization method added an additional term to the mean-squared-error cost function to limit deviations from the group-average model (see Methods), and thus allowed us to maintain the PK parameters within a physiologically reasonable range.

Figure 4A shows the six PK parameters in the QSP model that were randomly generated to reflect the variability in PK of durvalumab. The median capillary filtration rate was 5.6e-6 L/s, which is in agreement with our previous analysis of the relationship between experimentally observed permeability and the molecular size of durvalumab *(47)*. The median blood volume in the virtual patients is 5.8 liters, which agrees with the observed body weight distribution in the popPK study, given that the blood volume is approximately 70 mL/kg in adult humans. Further, the volume fraction of interstitial space in peripheral tissues available to durvalumab ranges from 3.5 to 7.9%, consistent with estimation in our previous study *(48)*. The clearance rates (both linear and non-linear) and Michaelis-Menten constant for non-linear clearance are also consistent with the estimated values in the popPK study *(46)*. The QSP model-predicted serum durvalumab concentration in the virtual patients is shown with clinically measured mean and standard deviation in Figure 4B. Consistent with clinical observations, durvalumab concentration reached steady state at approximately week 16 *(49)*. Model-predicted peak and trough concentrations after the first dose and the trough concentration at steady state were comparable to clinical measurements from Study 1108 and the ATLANTIC trial (Supplementary Table 1) *(49)*.

**Figure 4.**
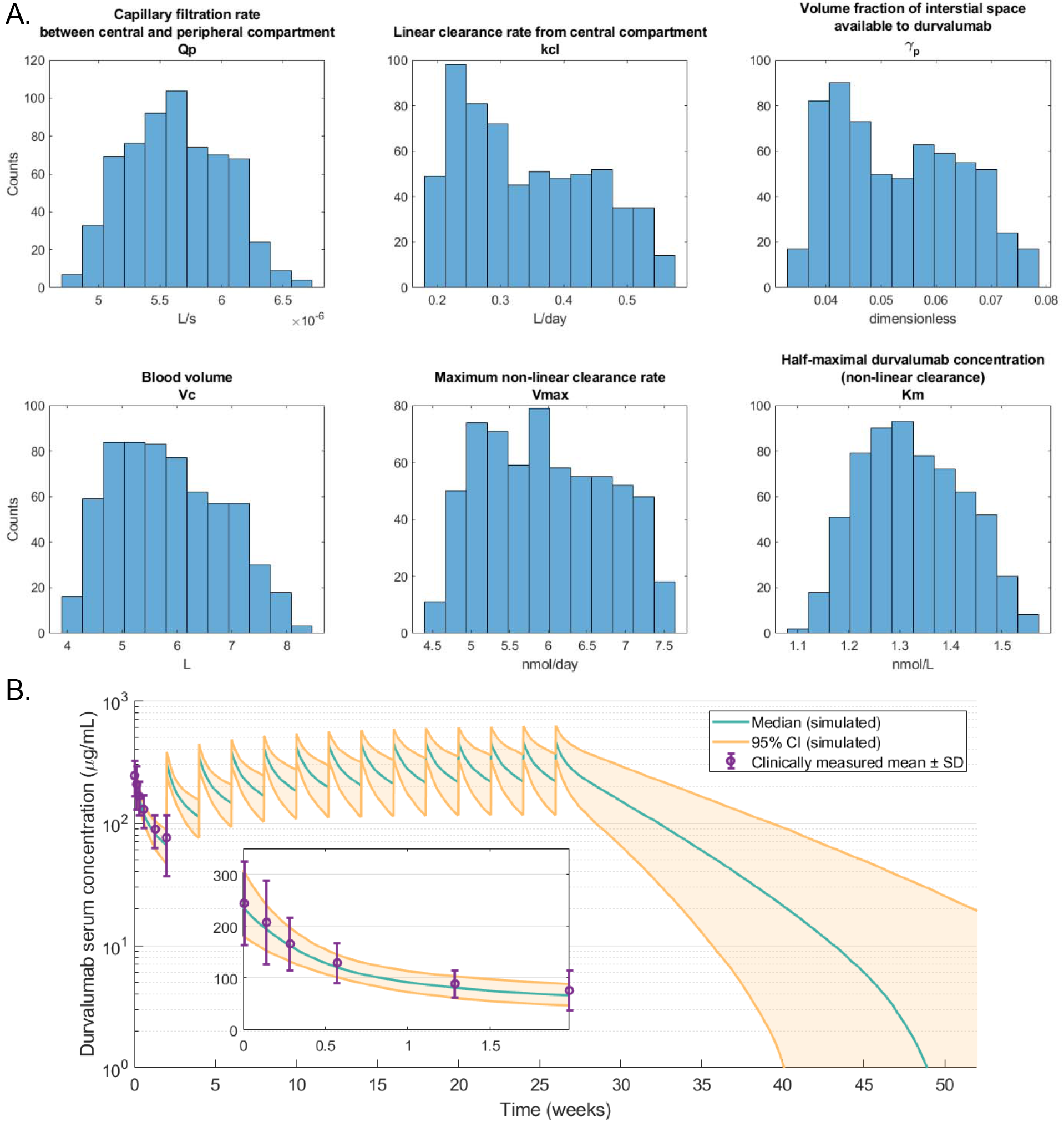
(A) Distribution of fitted pharmacokinetic parameter values in the quantitative systems pharmacology model and (B) model-predicted durvalumab serum concentration following a flat-dosing regimen of 750 mg every 2 weeks (Q2W). Green line represents the median model prediction; Orange lines represent the 5th and 95th percentiles.

### Predicting tumor dynamics during PD-L1 inhibition

With the PK parameters randomly generated from the latent space, we simulated PD-L1 inhibition in the virtual patient cohort. Durvalumab was administered with 750 mg flat doses every 2 weeks (Q2W) once each virtual patient reached the preset initial tumor diameter ranging from 1.5 to 9.9 cm (Figure 3). Tumor response to the treatment was analyzed in two virtual patient subgroups divided based on PD-L1 expression in the tumor with a threshold of 25%, which corresponds to the threshold used in Study 1108 *(24)*. In the clinical setting, PD-L1 expression is the percentage of tumor cells that exhibit membrane staining by the reagent *(24)*. According to our model prediction, the majority of the cells that express PD-L1 in the tumor were cancer cells, and since the PD-L1 density in the model was defined as the average PD-L1 level on cells in the tumor, we approximated the PD-L1 expression by dividing the model-predicted PD-L1 density by a theoretical maximum level of 1,770 molecules/μm^2^. The maximum level was estimated by the in vitro measurements of PD-L1 density on mature dendritic cells *(50, 51)*. Due to the lack of data on PD-L1 expression in NSCLC tumors, we estimated the average baseline PD-L1 expression (Table 1) so that the proportion of virtual patients that fell within each subgroup matched that reported by Study 1108 *(24)*.

Figure 5 shows the percentage change of tumor size and the best overall tumor size change in the two patient subgroups. Although the model predicted a faster median tumor growth for non-responders with a PD-L1-high tumor, responders in the PD-L1-high group showed a faster median tumor size reduction during early stage of the treatment. Specifically, 22.6% of the responders in the PD-L1-high group responded by week 6, as opposed to 6.1% in the PD-L1-low group. According to RECIST 1.1, the model predicted an objective response rate (ORR) of 18.6% with a 95% bootstrap confidence interval of (13.3, 24.2)% in the virtual patient cohort. In PD-L1-high and -low groups, ORRs were predicted to be 23.8 (16.3, 32.7)% and 12.0 (5.5, 20.2)%, respectively. The increase in ORR in the PD-L1-high group was potentially due to the positive correlation between PD-L1 expression and T cell infiltration (Supplementary Figure 2), which was also clinically observed *(43)*. Comparing to the model-predicted ORRs, the clinically reported ORRs in Study 1108 (21.8% and 6.4% for PD-L1 high vs. low) fall within the model-predicted 95% confidence intervals, while the difference in predicted ORRs between the two subgroups is narrower than that in the clinical trial. Our ORR prediction may be further improved by including new lesion formation in the model, since a large portion of the patients with low PD-L1 expression in Study 1108, despite having a >30% tumor size reduction, was categorized as non-responder due to detection of new lesion(s) *(24)*.

**Figure 5.**
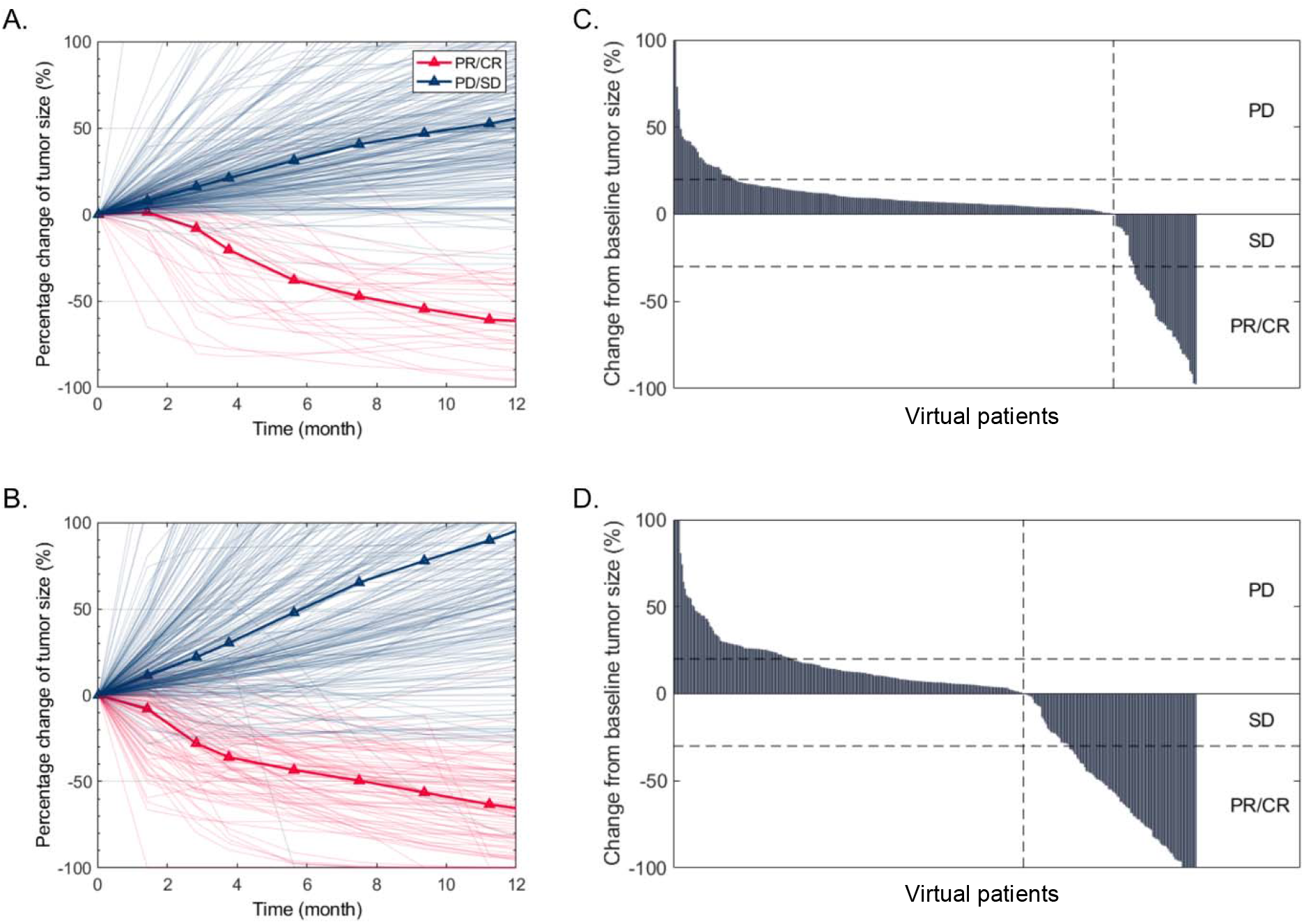
Percentage change in tumor size (A,B) and the best overall tumor size change (C,D) in PD-L1-low (A,C; N=276) and PD-L1-high (B,D; N=353) virtual patients during in silico clinical trials of durvalumab. R, responders; NR, non-responders; PD, progressive disease; SD stable disease; PR/CR, partial/complete response.

To further examine the performance of the virtual patient generation method, we simulated the same dose regimen of durvalumab in virtual patients selected by different combinations of immunogenomic data. Supplementary Table 2 shows that model-predicted ORRs were similar among virtual patient populations selected by any data combinations. This is likely because the parameter distributions that generate plausible patients were already manually calibrated to NSCLC data (see Methods). However, when we selected virtual patients by data on lung adenocarcinoma (LUAD) or lung squamous cell carcinoma (LUSD) separately, the model predicted higher ORR in LUSC with a median CD8/Treg ratio almost twice as high as that in LUAD, which aligned with clinical findings *(52)*. This observation suggests that the virtual patient generation method is capable of generating virtual patient populations that fit to particular patient subgroups while reducing the inherent uncertainty (as seen above in Figure 2A).

Next, we investigated the correlations between response status and PK variables, including the peak (*C*_*max*_), trough (*C*_*min*_) durvalumab concentration, and area under concentration curve (AUC) at early time points and steady state (week 16), as well as drug accumulation indices. Supplementary Figure 3 shows that the peak durvalumab concentration and AUC from day 0-14, as well as the peak concentration at steady state, were significantly higher in responders. We further divided the virtual patient into 5 subgroups with increasing level of each PK variable and calculated the ORR of each subgroup in Supplementary Figure 4. Interestingly, the ORR increased as *C*_*max*_ and AUC of the first dose (day 0-14) increased, with about a 14% difference in ORR between the 2 subgroups with the highest and the lowest level of *C*_*max*.1_ or *AUC*_0-14_. Nonetheless, a reverse trend, even though non-significant, was observed between ORR and *C*_*max*.1_ in patients with urothelial carcinoma from Study 1108 *(49)*.

To investigate the performance of potential predictive biomarkers in NSCLC, we compared their distributions between responders and non-responders from the overall virtual patient cohort. Figure 6 shows that responders have significantly higher CD8/4 T cells, PD-L1 level, CD8/Treg and CD8/CD4 ratios, number of NSCLC-specific T cell clones (TCC), and significantly lower MDSCs. Similar correlations between clinical response and CD8 T cell density, CD8/Treg and CD8/CD4 ratios were observed in a clinical trial of PD-1 inhibition in NSCLC *(43)*. In addition, the number of NSCLC-specific T cell clones was found to be correlated with tumor mutational burden *(53)*, which is a known predictive biomarker for immune checkpoint inhibition in NSCLC *(54)*. Next, we divided virtual patients into 5 subgroups with increasing level of each biomarker and calculated the ORR of each subgroup. Figure 7 confirmed that ORR increased as the biomarkers identified above increased (or decreased in case of MDSC). However, the disease control rate, which is defined as percentage of patients with complete/partial response and stable disease, increases only when CD8/Treg, CD8/CD4, and TCC increase or when MDSC decreases.

**Figure 6.**
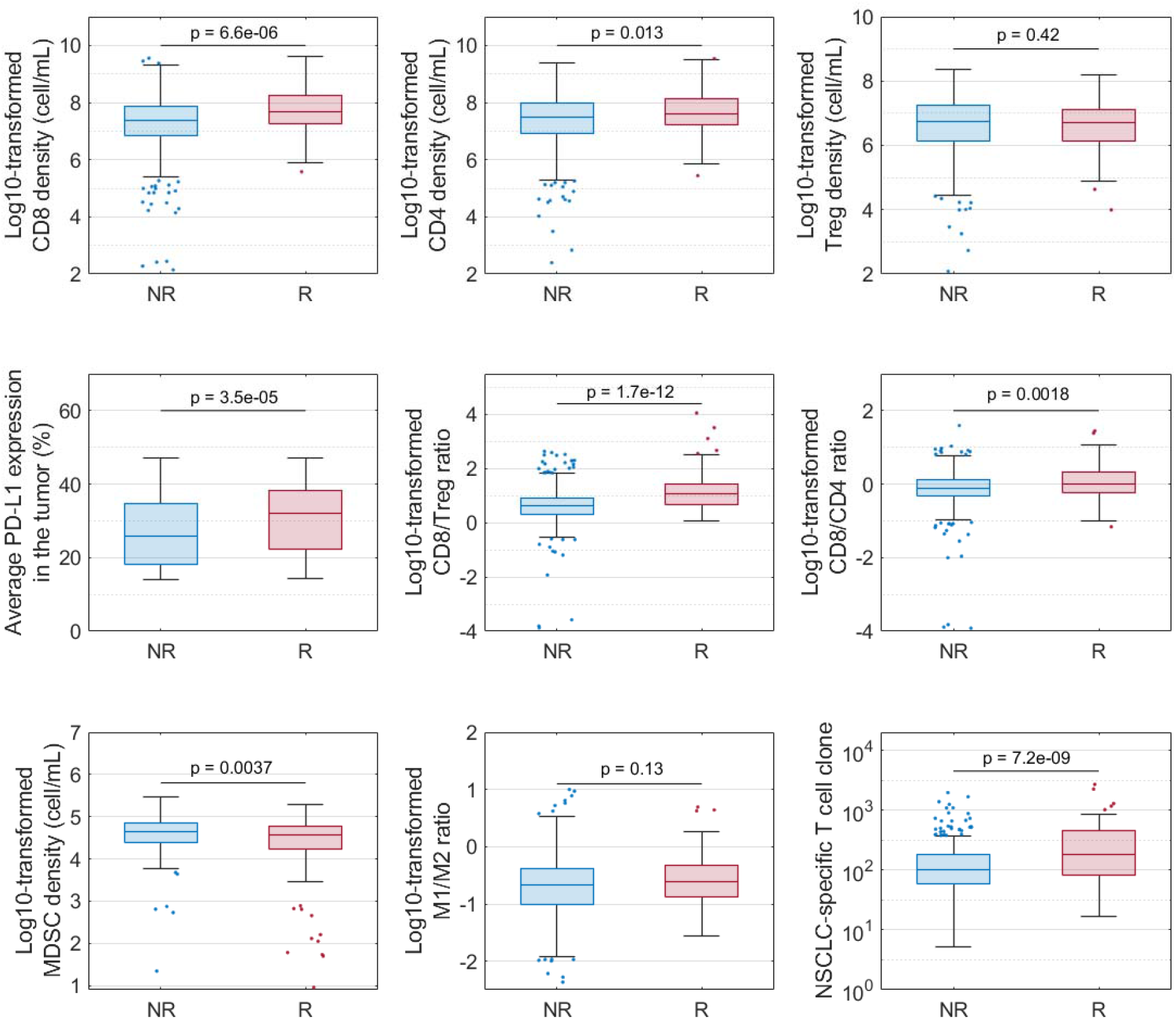
Comparison of pre-treatment variable distributions between responders (R) and non-responders (NR). p-values were calculated by Wilcoxon test.

**Figure 7.**
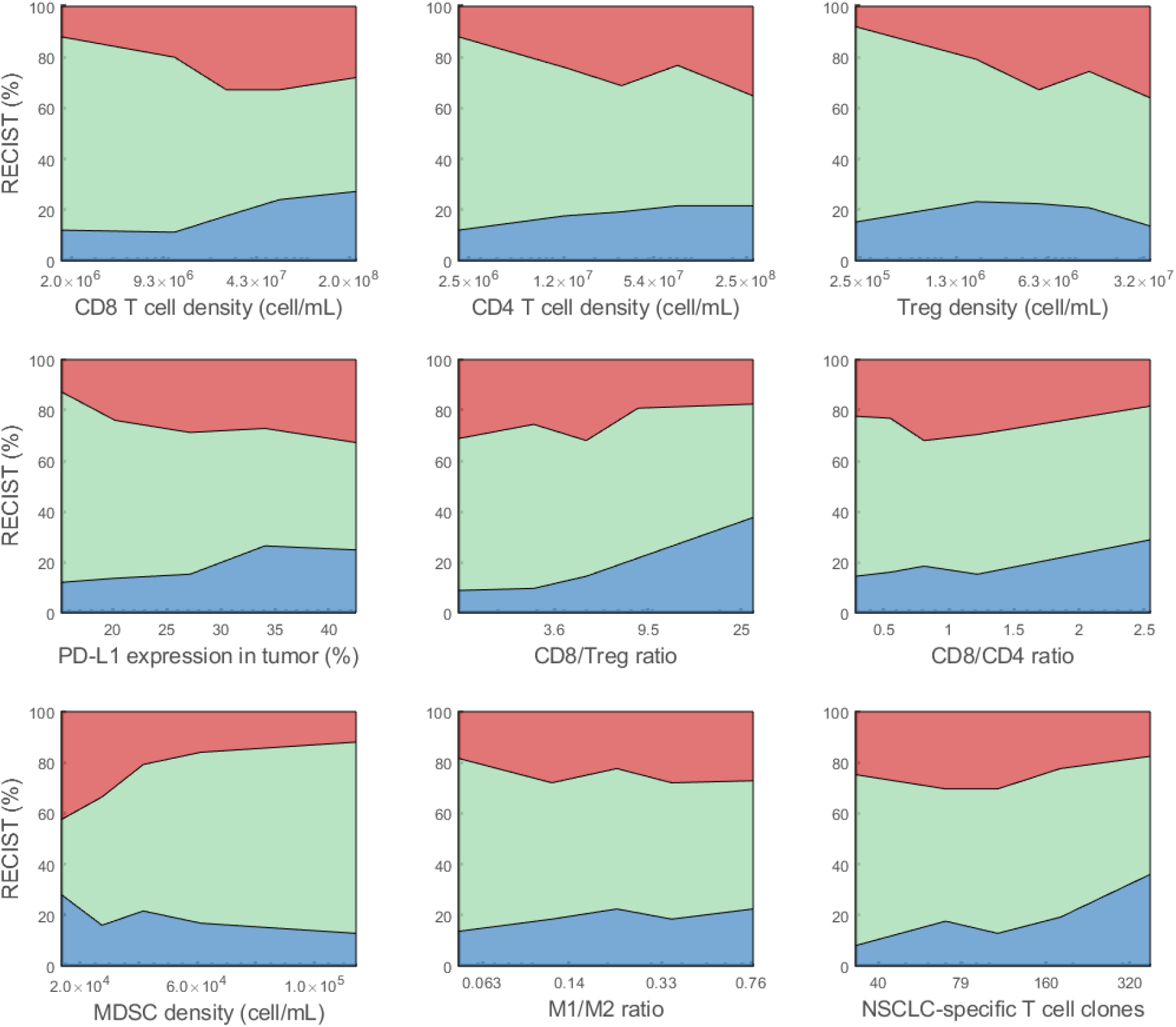
Effect of pre-treatment variables on objective response. For each variable of interest, virtual patients are sorted by the variable amount in ascending order, and evenly divided into 5 subgroups. The response status of each subgroup is plotted against the corresponding median variable amount. Blue represents partial or complete response. Green represents stable disease. Red represents progressive disease.

To study the combined effect of the predictive biomarkers, we trained a random forest model using pre-treatment CD8/4 T cells, PD-L1 expression, CD8/Treg, CD8/CD4 ratios, MDSC density, and TCC (see Methods). As a prerequisite, we calculated the correlation matrix to make sure that the variables were not strongly correlated (Supplementary Figure 5A). The variable importance of all pre-treatment biomarkers is shown in Supplementary Figure 6A, with CD8/Treg ratio, PD-L1 expression, CD8 density, and TCC identified as top important variables by the random forest model. With ROC analysis, we selected thresholds for the four predictive biomarkers that achieved a sensitivity of 80% (Figure 8A). The thresholds for CD8/Treg ratio, PD-L1 expression, CD8 density, and TCC were 4.1, 21%, 151 cells/mm^2^, and 72 (with specificity of 47%, 35%, 37%, and 34%), respectively. Here, we converted the CD8 density from 3-D to 2-D that corresponds to the outcome of digital pathology analysis *(26)*. Further, we performed similar analyses on on-treatment biomarkers. We selected CD8/Treg ratio and CD8 density at the end of first treatment cycle (day 14), which were identified by the random forest model as the two most important variables (Supplementary Figure 5B and 6B). The thresholds, according to ROC analysis in Figure 8B, were selected to be 5.1 and 176 cells/mm^2^ (with specificity of 52% and 41%). Overall, CD8/Treg ratio showed the best performance with the highest specificity and area under ROC curve (Figure 8). In comparison, Kim et al. analyzed tumor-infiltrating lymphocytes in 33 primary lung lesions from advanced NSCLC and found that a Treg/CD8 ratio cutoff of 0.25 achieved a sensitivity and specificity of 82.6% and 65.4% in predicting response to anti-PD-1 treatment *(43)*. In a meta-analysis, Li et al. also identified tumor PD-L1 expression and mutational burden as predictive biomarkers (both with specificity of about 30% when sensitivity is 80%) for anti-PD-(L)1 treatment in NSCLC *(55, 56)*.

**Figure 8.**
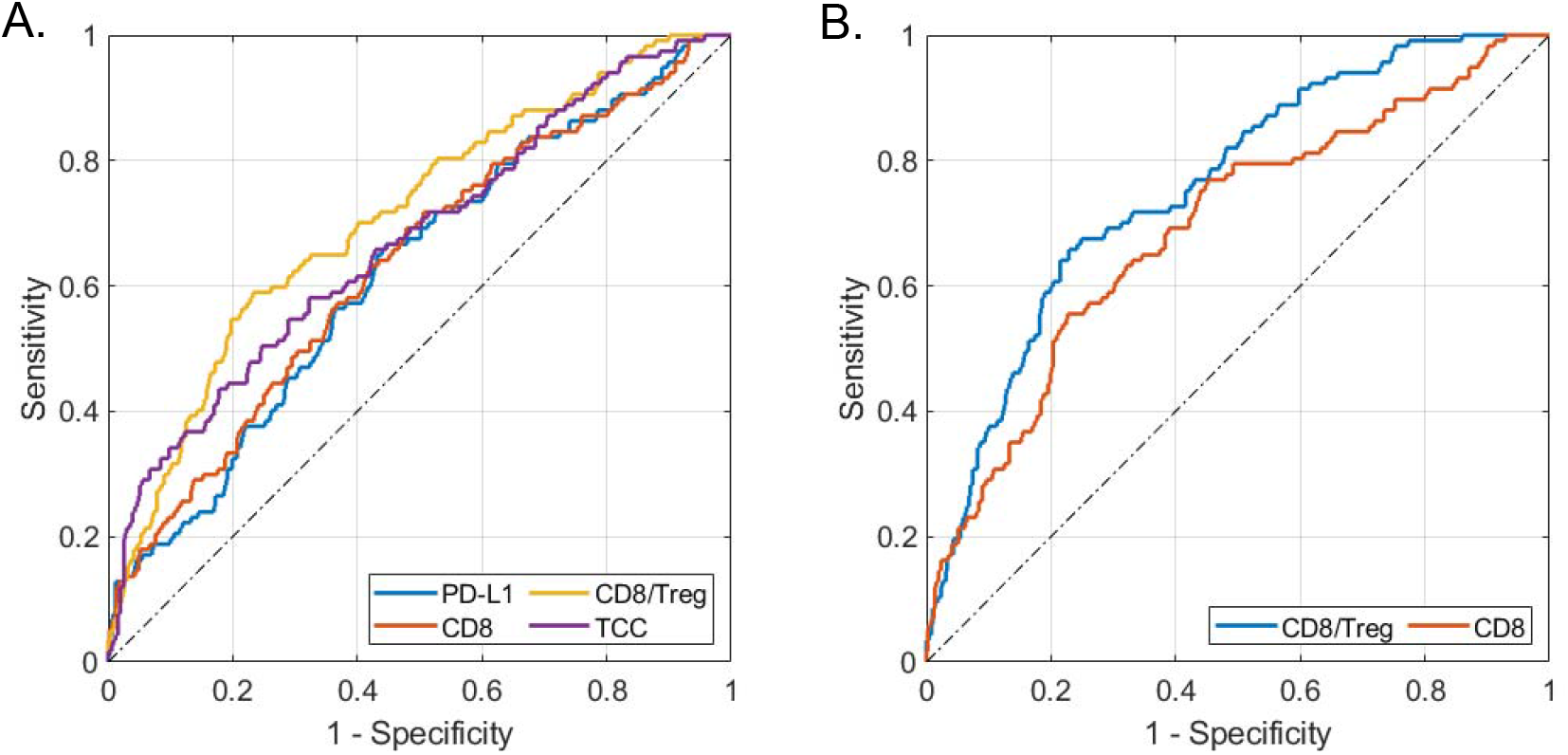
ROC analysis of (A) pre-treatment and (B) on-treatment predictive biomarkers for PD-L1 inhibition in NSCLC. Areas under curve (auROC) were 0.62, 0.63, 0.71, 0.67 for pre-treatment PD-L1 expression, CD8 T cell density, CD8/Treg ratio, and NSCLC-specific T cell clones (TCC), respectively; auROC were 0.76 and 0.69 for on-treatment CD8/Treg ratio and CD8 density at the end of first treatment cycle (day 14).

To explore dynamics of immune cells during immunotherapy, we visualized the time-dependent profiles of immune cells in the central (Supplementary Figure 7) and tumor compartment (Supplementary Figure 8). Supplementary Figure 7 shows that all activated T cell subsets in blood, including CD8, CD4, helper T cells, and Tregs were increased by durvalumab in responders, while CD8/Treg and CD8/CD4 ratios decreased over time in responders. On the contrary, immune cells in the non-responders had opposite dynamics when compared to the responders. For immune cells in the tumor, Supplementary Figure 8 shows that activated T cell subsets also increased in responders during PD-L1 inhibition. However, unlike T cell ratios in blood, CD8/CD4 ratio increased in responders, and CD8/Treg ratio was transiently increased by durvalumab in responders for the first two months and did not drastically change from the baseline level in the long term. Furthermore, M1/M2 ratio decreased and MDSC density increased in responders, suggesting that TAMs and MDSC may partly contribute to resistance to immunotherapy in NSCLC *(57)*. Sensitivity analysis (Supplementary Figure 9) also suggested that recruitment of MDSC, Th-to-Treg differentiation, and M2 polarization could be potential targets of drugs that can be combined with durvalumab, as parameters related to these mechanisms were among the most influential ones to tumor size at the end of durvalumab treatment.

## DISCUSSION

In the present study, we revisited PD-(L)1 simulation in NSCLC with the latest QSP model expansion and attempted to address the challenge on generating heterogeneous yet physiologically realistic virtual patients, which was raised in our previous studies *(13–15)*. In terms of model structure, we utilized previously developed QSP model of TNBC and incorporated an additional source of IFNγ in the tumor microenvironment (see Methods). In addition, baseline values of cancer type-specific parameters were recalibrated by experimental and clinical data on stage III NSCLC. Notably, PD-L1 in the model represents the average expression on all tumor cells, including cancer cells and immune cells, that can interact with PD-1 on activated T cells and TAMs to inhibit Teff-mediated cancer killing and TAM-mediated phagocytosis. However, PD-L1 on different cell types may have different roles in the tumor microenvironment *(58)*, and thus can be separately modeled in future studies. In addition, PD-L1 expression is upregulated only by IFNγ in the current model, so we assume a baseline PD-L1 expression to match the percentage of virtual patients in the PD-L1-high and PD-L1-low groups as observed in the clinical trial. It should be noted that multiple inflammatory signaling pathways are involved in PD-L1 upregulation, which results in the clinically observed heterogeneous PD-L1 expression *(59)*. Additional mechanistic details can be incorporated into the model when sufficient experimental data become available for model calibration.

One of our main focuses is to improve our approach to virtual patient generation. In previous studies, we randomly generated deviations around baseline parameter values assuming uniform, log-uniform, or log-normal distribution, and manually calibrated the distribution statistics (e.g., standard deviation, upper and lower boundaries) so that the summary statistics in virtual patients matched those reported by clinical analyses *(15)*. This is a time-consuming process, which becomes even more challenging when fitting to multi-dimensional data. The probability of inclusion proposed by Allen et al. allowed us to select virtual patients that statistically matched the observed data from the randomly generated plausible patients *(21)*, and iAtlas portal provided cancer-specific patient-level data for this method. However, unlike the model of cholesterol metabolism presented by Allen et al., physiologically plausible ranges for the present QSP model variables are not well established due to the insufficient biological understanding of the tumor microenvironment. Therefore, we could not apply the additional step to optimize plausible patients via simulated annealing before calculating the probability of inclusion, which would further improve confidence in model predictions by constraining virtual patients within the physiologically reasonable ranges *(21)*. Comparing to the results from Allen et al., similar proportion (2-3%) of plausible patients in this study was included in the final virtual patient cohort, which depended on the data dimensionality and the initial distribution of plausible patients *(21, 60)*.

Besides the three immune subset ratios that were used to select virtual patients in the present analysis (Figure 2), there are other patient characteristics, such as cancer cell growth rate, T cell clonality, and binding affinity between neoantigen and MHC molecules, which also differ among patients but cannot be directly obtained from the immunogenomic data. In future analyses, machine learning algorithms can be applied on multi-omics data to predict model-related parameter values *(61–63)*. Importantly, we have demonstrated here that the virtual patients generated by the QSP model and selected by the three immune subset ratios were able to capture the inter-patient heterogeneity while being consistent with unseen digital pathology data on immune cell densities in NSCLC tumor.

To account for variability in PK of durvalumab and predict its intratumoral concentration, we fitted PK parameters in the QSP model to the data simulated from the time-dependent popPK model *(46)* via compressed latent parameterization. This method not only considers the covariance between parameters but also minimizes parameter deviations from the group-average value when there is a lack of parameter identifiability *(23)*. One of the limitations, however, is that the popPK data and the immunogenomic data were collected from two different patient populations, so the covariances between PK-related characteristics and immunogenomic data were not accounted for during virtual patient generation. In addition, as the patient data in iAtlas portal were not body weight dependent, we simulated 750 mg flat dosing instead of weight-based dosing.

With the virtual patients that have shown resemblance to real patient data, we investigated if the model could make reliable efficacy prediction for durvalumab in stage III NSCLC. Following the same dose regimen in Study 1108, we simulated 750 mg doses every 2 weeks and divided virtual patients into two subgroups based on their PD-L1 expression on tumor cells. The model predictions fell within the clinically reported confidence intervals and confirmed that patients with high PD-L1 expression have a higher ORR than the PD-L1-low group. It should be noted that appearance of new lesions (e.g., locoregional and distant metastases) is not a rare event especially during treatment of late-stage NSCLC *(24, 64)*, which would be categorized as progressive disease regardless of the tumor size change. New lesions are most likely seeded before therapy begins and grow to a detectable size during the treatment *(65, 66)*. This stochastic process, which may require a hybrid modeling technique or approximation methods to integrate into the QSP model *(67, 68)*, may be addressed if relevant data become available in the future.

## MATERIALS AND METHODS

### Overview of the QSP modeling platform for immuno-oncology (QSP-IO)

The QSP model comprises four main compartments: central (C), peripheral (P), tumor (T), and tumor-draining lymph node (LN). These compartments represent the circulating blood, lumped peripheral tissues/organs, tumor microenvironment, and lumped tumor-draining lymph nodes, respectively. Ten modules were involved in this study, which described dynamics of cancer cells, T cells (i.e., effector, helper, and regulatory T cells), antigen-presenting cells, neo/self-antigens, therapeutic agent, immune checkpoints, myeloid-derived suppressor cells (MDSCs), and tumor-associated macrophages (TAMs). Cancer type-specific parameters were reparametrized to NSCLC based on experimental data, preferentially from stage III NSCLC if available (Table 1). In addition, we modified our previous assumption so that the main source of interferon-gamma (IFNγ) was not limited to activated CD4 helper T cells. Instead, we assumed that activated CD8 T cells also produced IFNγ with a rate 3 times higher than CD4 T cells *(69)*. For cancer cell growth, we assumed logistic growth for NSCLC with a constant maximum carry capacity of 10 cm (Table 1). Other mechanisms remained the same as in our previous analysis, which was elaborated in *(15)*.

Overall, there are 255 parameters, 141 ODEs, and 40 algebraic equations (model rules with repeated assignments). Model simulations were performed using SimBiology Toolbox in MATLAB R2020b (Mathworks, Natick, MA) with ODE solver, SUNDIALS. Each simulation started from a single cancer cell, and tumor volume was calculated at each time step via Equation 1. *C*_*total*_, *T*_*total*_, and *M*_*total*_ are the total number of cancer cells, T cells, and TAMs; *V*_*cell*_, *V*_*Tcell*_, and *V*_*Mcell*_ are volumes of single cancer cell, T cell, and macrophage; *C*_*x*_ and *T*_*exh*_ are the numbers of dying cancer cells and exhausted T cells; and *V*_*e,T*_ is the volume fraction of intracellular space in NSCLC tumor. During postprocessing, tumor diameter was estimated assuming a spherical tumor.

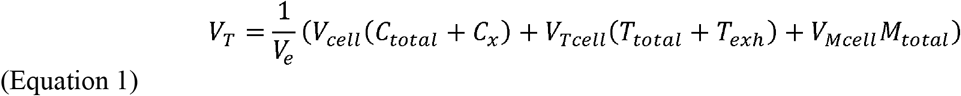

### Virtual patient generation

To account for interindividual variability that results in the heterogeneous tumor response to immunotherapy, we selected 30 out of the 255 model parameters, which are known to differ among patients, and generated random deviations around their baseline values using Latin-hypercube sampling (LHS). The distribution statistics (e.g., standard deviation, upper and lower boundaries) were estimated based on available clinical observations and experimental data on NSCLC. In theory, each randomly generated parameter set represents a virtual patient *(70)*. However, to avoid confusion, we define the virtual patients generated at this step as plausible patients. Notably, each patient was randomly assigned a preset initial tumor diameter (see *initial_tumour_size* in Table 1) *(40, 41)*. When the tumor size reached the preset value, the model variables were saved and treated as the pre-treatment characteristics for the corresponding patient. At this step, we generated 30,000 plausible patients.

To generate a virtual patient cohort whose characteristics statistically match the real patient population, we adapted the probability of inclusion proposed by Allen et al. *(21)*. We first explored the “Immune Cell Proportions” data from the “Immune Feature Trends” module in iAtlas portal (https://isb-cgc.shinyapps.io/shiny-iatlas/) by selecting TGCA subtypes: lung adenocarcinoma (LUAD) and lung squamous cell carcinoma (LUSC), which are the two major subtypes of NSCLC *(22, 71)*. Then, we downloaded the proportions of CD8, CD4, and regulatory T cell (Treg), M1 and M2 macrophages from analyses of the two TCGA subtypes; and we generated a 3-dimensional patient-level data containing ratios between CD8 T cells and Tregs, CD8 and CD4 T cells, and M1 and M2 macrophages, all of which have corresponding QSP model species. Data points that contained zero value(s) were removed to avoid singularities. Moreover, as the TCGA data were collected from untreated patients, they correspond to the pre-treatment characteristics predicted by the model *(72)*. Finally, we calculated the probability of inclusion via Equation 2, which is the conditional probability of including a plausible patient into the final virtual patient cohort (i.e., *S*(*θ*) = 1) given the model prediction *M*(*θ*) = *r*.

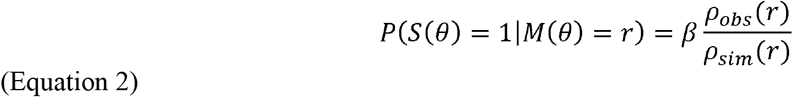

Here, *S* is a logical function that equals to 1 if the plausible patient *θ* should be included and 0 if otherwise, and *M* represents the QSP model that predicts time-dependent profile of model variables for the plausible patient *θ*. In this study, we focused on the model-predicted pre-treatment ratios that corresponded to the data from iAtlas (i.e., CD8/Treg, CD8/CD4, and M1/M2). Additionally, *ρ*_*obs*_(*r*) and *ρ*_*stm*_(*r*) are the probability density estimate for model-predicted ratios, *r*, in the observed data and the plausible patients, respectively. Probability densities were estimated by 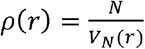, where *V*_*N*_(*r*) is the volume of an N-dimensional hypersphere with radius defined by the distance to the N-th nearest-neighbor of *r. N* was typically chosen from 5 to 10 in this study. The constant *β* was optimized by the simulated annealing algorithm to minimize the average two-sample Kolmogorov-Smirnov test statistic when evaluating the difference between the empirical cumulative distribution functions of the observed data from real patients and the predicted values from selected virtual patients *(21)*. With the optimal *β*, virtual patients were selected based on the inclusion probability to generate the final virtual patient cohort.

### In silico clinical trial

To account for the interindividual variability in pharmacokinetics of durvalumab, we applied compressed latent parameterization proposed by Tivay et al. *(23)* to generate PK parameters for the virtual patients. Since this method required patient-level PK data for durvalumab, which were not available from published studies, we first generated time-dependent durvalumab PK of 400 pseudo-patients based on the population PK (popPK) study by Baverel et al. *(46)*. Specifically, we assumed log-normal distribution for patient characteristics, such as serum albumin level, body weight, and soluble PD-L1 level, and we estimated the standard deviations based on the reported means and ranges. For categorical variables, including sex and Eastern Cooperative Oncology Group (ECOG) performance status, we assumed binomial distribution with probabilities estimated by the summary statistics of corresponding patient characteristics. Based on the estimated distributions, we randomly generated characteristics of 400 pseudo-patients and calculated values for linear clearance, *cl*, maximum elimination rate, *V*_*max*_, and volumes of the two compartments, *V*_1_ and *V*_2_, in the popPK model using algebraic equations provided by Baverel et al. *(46)*. Intercompartmental clearance, *Q*, was assumed to be 0.476 L/day for all patients. Since time-dependent profile of the patient characteristics were not available, we assumed they were constant over time. With the following ODEs for the two-compartment popPK model, we simulated the durvalumab concentration profile in the 400 pseudo-patients *(46)*.

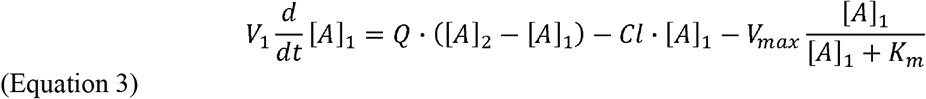

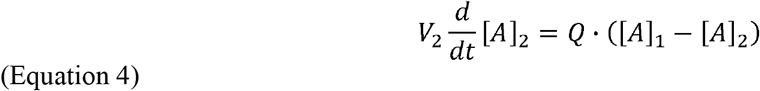

The first step of compressed latent parameterization is to find the “group-average” model defined by 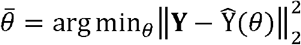, where **Y** is a matrix storing data from all patients and 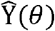 is the corresponding model predictions *(23)*. Here, due to the high variability in durvalumab PK, we defined **Y** as an Nt-by-1 vector storing the median durvalumab concentration in the 400 pseudo-patients at each time point (Nt is the length of the time vector), and 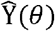 as an another Nt-by-1 vector storing the QSP model-predicted durvalumab concentration given an input PK parameter set *θ*. We fitted six PK parameters, which were capillary filtration rate, blood volume, volume fraction of interstitial space in peripheral tissues available to durvalumab, linear and maximum non-linear clearance rates, and Michaelis-Menten constant for non-linear clearance, using MATLAB function, fmincon. Then, we randomly generated k=500 random local deviations around the group-average model, which was stored in a 6-by-k matrix **Θ**. The corresponding changes in model-predicted serum durvalumab concentration were stored in a Nt-by-k vector 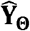. Through singular value decomposition of the covariance matrix 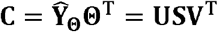, we got a 6-by-6 matrix **V** whose columns are sorted orthogonal directions of maximum covariance in the parameter space *(23)*. Further, we fitted the six PK parameters in the QSP model to the pseudo-patient data generated above via 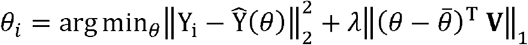, and a latent parameter space was constructed via 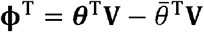, where *θ* is a 6-by-400 matrix storing all fitted parameter sets. Finally, new PK parameter sets were randomly generated from the latent space via 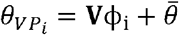 for the virtual patients, where the 6-by-1 vector ϕ_i_ was randomly sampled from the latent space **ϕ** assuming independent uniform distribution for each dimension of the latent parameter space.

With the final virtual patient cohort, we simulated their tumor response to durvalumab treatment starting from the time point when the tumors reached their preset initial diameter (i.e., pre-treatment tumor size), following the same settings in a phase 1/2 clinical trial of durvalumab (NCT01693562) *(24)*. 10 mg/kg durvalumab was administered every 2 weeks (Q2W) via a SimBiology dose object. Tumor diameters were recorded at 6, 12, and 16 weeks, and every 8 weeks thereafter, corresponding to the frequency of pre-scheduled tumor size measurement in the clinical trial. Clinical response was classified by RECIST v1.1 *(73)* with a minimum duration of stable disease of 6 weeks.

### Statistical analyses

For comparison between model-predicted ORRs and clinical observation, bootstrap sampling was performed to resample the virtual patient (sub)population with a sample size matching the number of patients in the corresponding subgroups (i.e., PD-L1 high/low) in Study 1108. The bootstrap median and the 95 percentile confidence intervals were then calculated. Wilcoxon tests were conducted using ranksum function in MATLAB 2020b.

Random forest models were trained on potential predictive biomarkers of interest to predict response status using caret and randomForest packages in R 4.2.3. For each model, 500 trees were trained, and each tree was trained on two-thirds of the data points. The out-of-bag error is defined as the error rate of each tree in predicting the data excluded by the training set (i.e., out-of-bag samples). The Mean Decrease Accuracy for each variable is the average decrease of model accuracy in predicting outcomes of the out-of-bag samples when a particular variable is excluded from the model, which is reported as variable importance. Receiver Operating Characteristic (ROC) analyses were performed by perfcurve function in MATLAB 2020b.

Sensitivity analysis was performed using Morris screening/elementary effects method *(74, 75)* with a 11-level grid and a step size Δ of 1/10 for the 30-dimensional hypercube [0,1]^30^ (i.e., 30 input parameters varied during virtual patient generation). Actual values of the 30 parameters were calculated based on their distributions by treating the sampled points from the hypercube as quantiles *(75)*. 1000 trajectories were randomly generated at the beginning. Trajectories with points fell outside the hypercube, as well as those with points containing 0% and 100% quantiles for parameters with lognormal distribution, were disregarded. Overall, 34 successful trajectories were plugged into the model to simulate tumor sizes at day 400 of durvalumab treatment, which were used to calculate the elementary effects (EEs). Finally, the variance *σ*^2^ and the mean absolute values *μ*^*^ of the EEs were estimated for each parameter *(75)*.

## Acknowledgments

The authors thank Thomas K. Kilvaer for sharing their data from digital pathology analysis; and lab members Alberto Ippolito, Shuming Zhang, Babita Verma, Samira Anbari, and Mehdi Nikfar for reading the manuscript and critical comments. The authors also thank Babita Verma for advice on statistical analysis.

## Author contributions

Conceptualization: A.S.P.; Methodology: H.W.; Resources: T.A., H.W.; Investigation: H.W.; Visualization: H.W.; Funding acquisition: A.S.P.; Supervision: A.S.P.; Writing – original draft: H.W.; Writing – review & editing: H.W., T.A., A.S.P., H.K.

