## Supplementary for "Generating immunogenomic data-guided virtual patients using a QSP model to predict response of advanced NSCLC to PD-L1 inhibition"

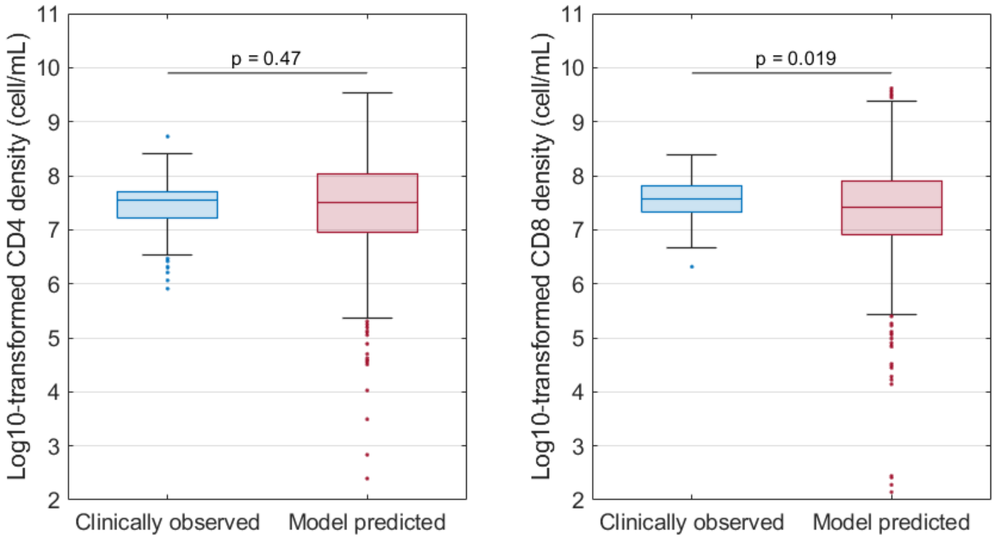


**Supplementary Figure 1. Comparison of CD8 and CD4 T cell density distributions between virtual cohort and patient population with stage III NSCLC requested from Kilvaer et al. (PMID: 33035322).** p-values were calculated by Wilcoxon test.

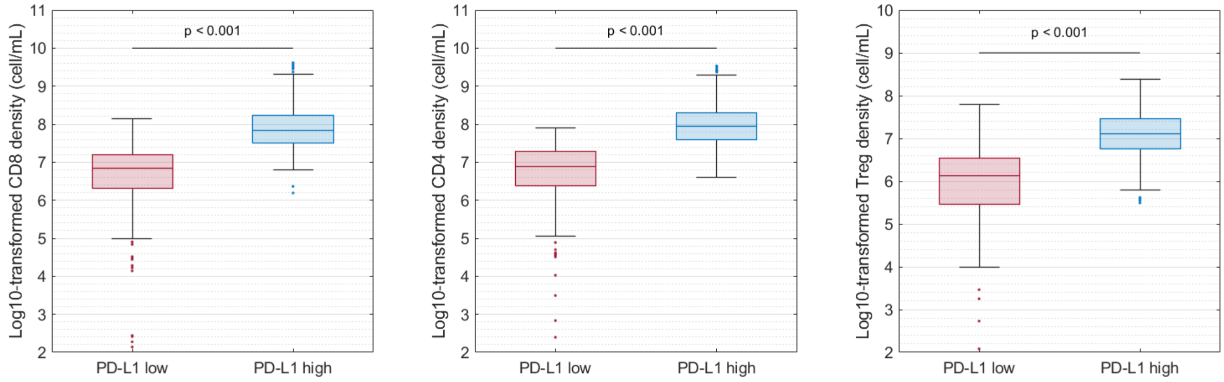


**Supplementary Figure 2. Comparison of pre-treatment immune cell densities between PD-L1-low and PD-L1-high groups.** p-values were calculated by Wilcoxon test.

**Supplementary Table 1. Comparison between model-predicted pharmacokinetics and clinical observation.**

| | Cmax,1 (µg/mL) | Cmin,2 (µg/mL) | Cmin,ss (Ctrough,w16; µg/mL) |
|---|---|---|---|
| Model prediction[a] | 236 (207, 271; N=629) | 66.2 (56.6, 75.9; N=629) | 209 (149, 275; N=629) |
| Clinical measurement (Study 1108)[b] | 222 (185, 260; N=838) | 81.4 (56.6, 112; N=819) | 162 (121, 217; N=334) |
| Clinical measurement (ATLANTIC trial)[c] | 204 (173, 229; N=399) | 86.9 (68.9, 107; N=355) | 165 (116, 204; N=193) |

Median and (25, 75) percentiles are listed with the number of patients/virtual patients. Cmax,1 is the post-first dose maximum durvalumab plasma concentration. Cmin,2 is the pre-second dose minimum concentration. Cmin,ss is the minimum/trough concentrations at steady state (week 16).

[a] US Food and Drug Administration. Clinical pharmacology and biopharmaceutics review(s) for Application number 761069Orig1s000. https://www.accessdata.fda.gov/drugsatfda_docs/nda/2017/761069Orig1s000ClinPharmR.pdf

[b] NCT01693562

[c] NCT02087423

**Supplementary Table 2. Model-predicted objective response rate (ORR) and immune subset ratios in virtual patients selected by various data combinations.**

| | Predicted ORR (95% CI) | CD8/Treg (95% CI) | CD8/CD4 (95% CI) | M1/M2 (95% CI) |
|---|---|---|---|---|
| Immunogenomic data[a] | N/A | 4.9 (0.9, 121.9) | 0.9 (0.2, 10.9) | 0.22 (0.01, 0.90) |
| Plausible patients | 16.8 (11.3, 22.3)% | 5.1 (0.6, 87.7) | 1.1 (0.2, 8.8) | 0.31 (0.03, 5.13) |
| VP filtered by M1/M2 | 16.6 (11.3, 22.3)% | 5.4 (0.6, 150.6) | 0.8 (0.1, 6.2) | 0.24 (0.02, 2.34) |
| VP filtered by CD8/Treg | 16.1 (10.6, 21.9)% | 5.0 (0.7,114.1) | 0.8 (0.1, 5.1) | 0.22 (0.02, 1.88) |
| VP filtered by CD8/CD4 | 18.6 (13.3, 24.2)% | 5.3 (0.7, 112.6) | 0.8 (0.1, 5.7) | 0.24 (0.02, 2.37) |
| VP filtered by M1/M2 and CD8/Treg | 15.9 (10.9, 21.1)% | 4.9 (0.6, 108.5) | 0.8 (0.1, 7.3) | 0.22 (0.02, 2.25) |
| VP filtered by M1/M2 and CD8/CD4 | 18.8 (13.7, 24.6)% | 5.4 (0.6, 99.3) | 0.8 (0.1, 6.1) | 0.24 (0.02, 1.97) |
| VP filtered by CD8/Treg and CD8/CD4 | 17.80 (12.70, 23.83)% | 5.3 (0.7, 140.7) | 0.8 (0.1, 6.3) | 0.24 (0.02, 3.23) |
| VP filtered by LUSC | 20.6 (14.5, 27.0)% | 7.0 (0.8, 159.6) | 1.1 (0.1, 9.8) | 0.26 (0.02, 2.73) |
| VP filtered by LUAD | 16.5 (11.1, 22.7)% | 4.0 (0.7, 79.9) | 0.7 (0.1, 4.9) | 0.20 (0.02, 3.23) |
| Virtual patients | 18.6 (13.3, 24.2)% | 5.0 (0.5, 178.3) | 0.8 (0.1, 6.5) | 0.22 (0.02, 2.53) |

[a] Data points that contained zero value(s) for immune cell proportions were removed to avoid singularities.

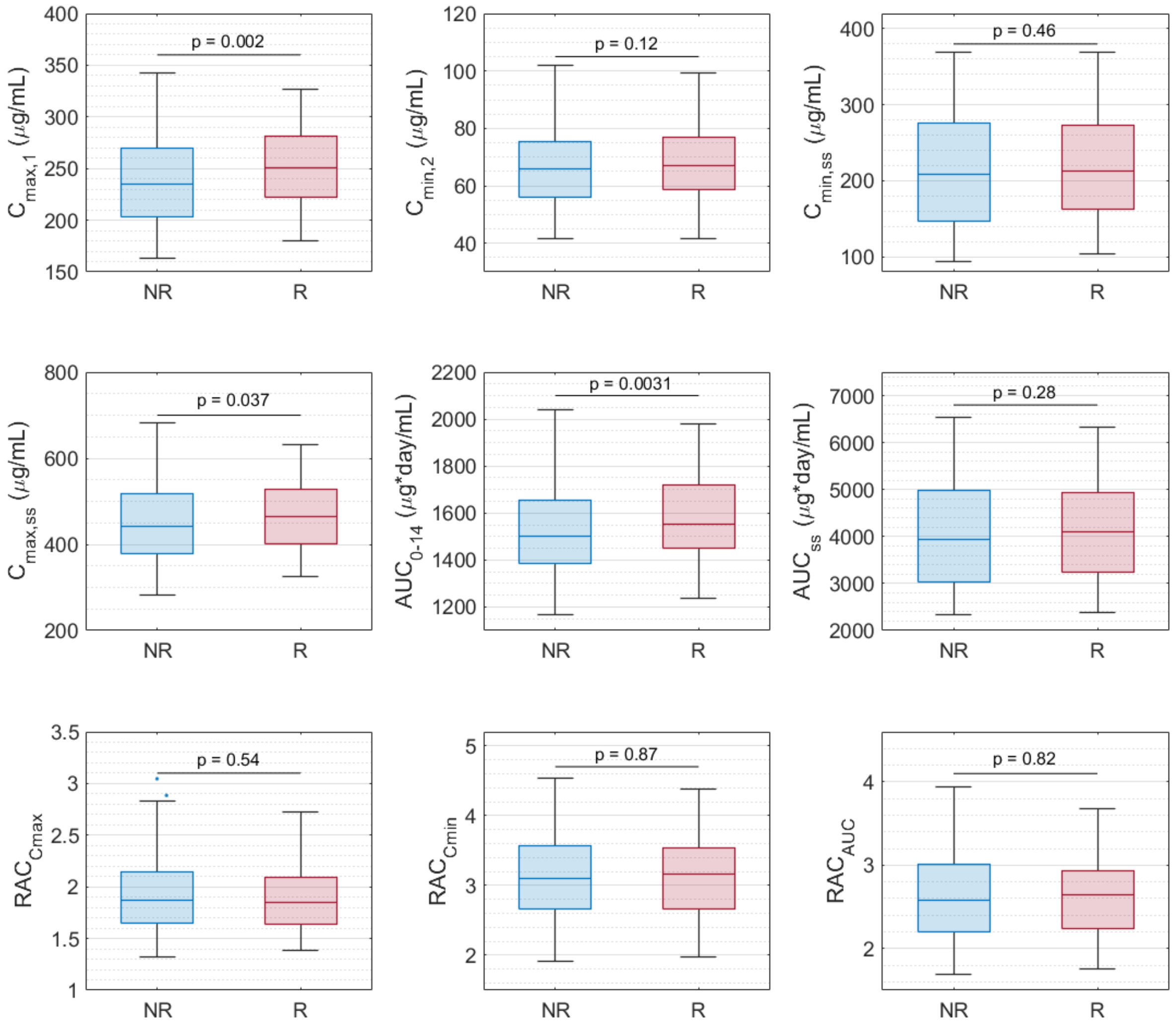


**Supplementary Figure 3. Comparison of pharmacokinetic variable distributions between responders (R) and non-responders (NR).** Cmax,1 is the post-first dose maximum durvalumab plasma concentration. Cmin,2 is the pre-second dose minimum concentration. Cmax,ss and Cmin,ss are the maximum and minimum concentrations at steady state (week 16), respectively. AUC0-14 is the area under the concentration curve from day 0-14. RACs are the drug accumulation ratios of Cmax,ss/Cmax,1, Cmin,ss/Cmin,2, and AUCss/AUC0-14. p-values were calculated by Wilcoxon test.

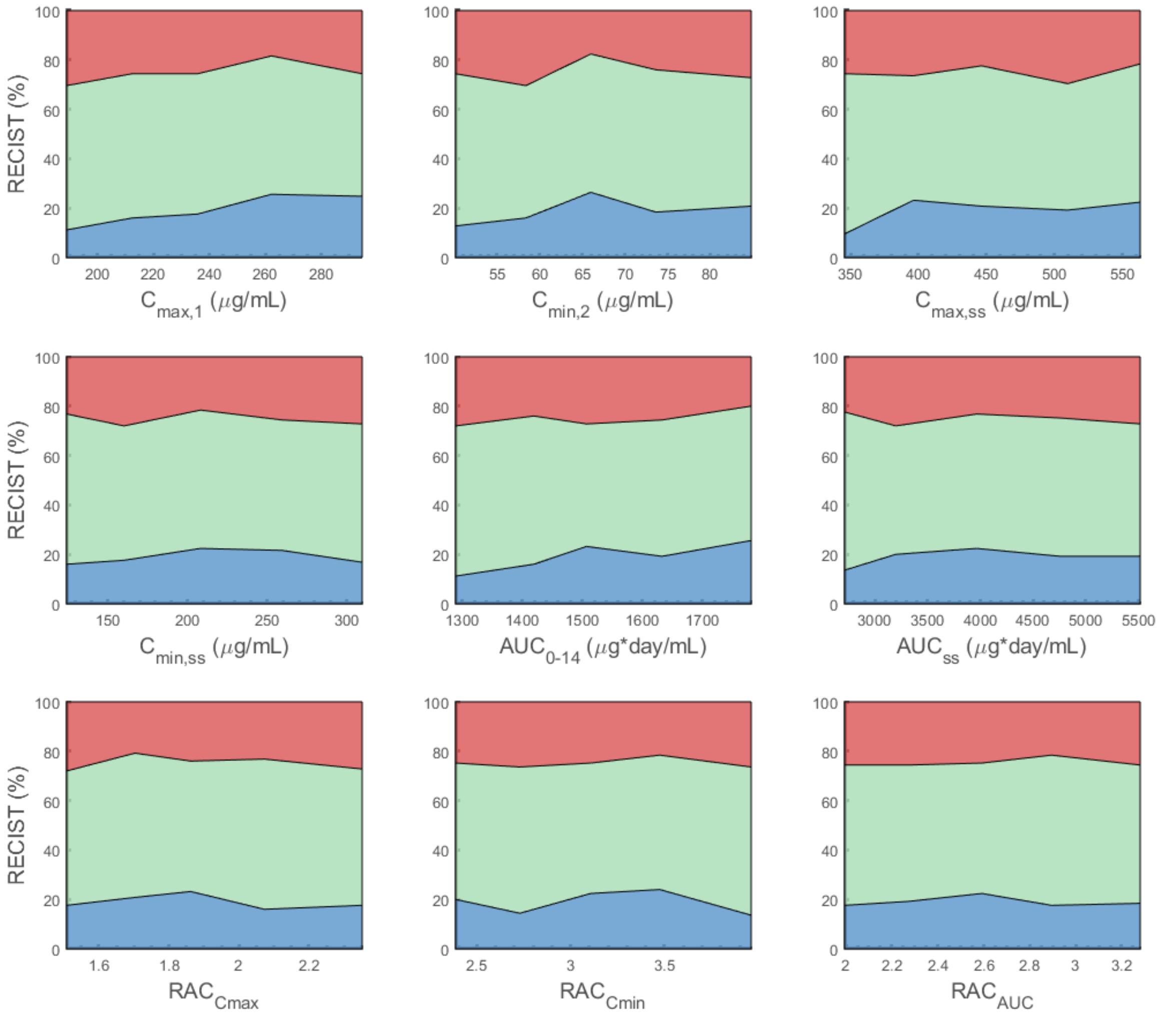


**Supplementary Figure 4. Effect of pharmacokinetic variables on objective response.** For each variable of interest, virtual patients are sorted by the variable amount in ascending order, and evenly divided into 5 subgroups. The response status of each subgroup is plotted against the corresponding median variable amount. Blue represents partial or complete response. Green represents stable disease. Red represents progressive disease.

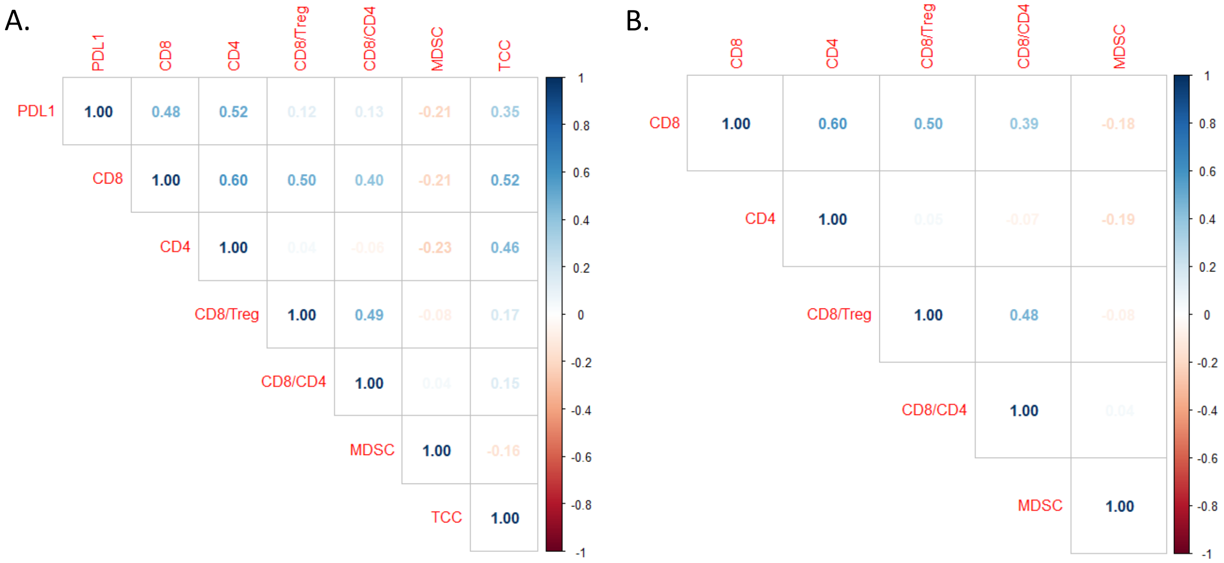


**Supplementary Figure 5. Correlation matrix of (A) pre-treatment and (B) on-treatment variables.** Treg, regulatory T cell. MDSC, myeloid-derived suppressor cells. TCC, the number of NSCLC-specific T cell clones.

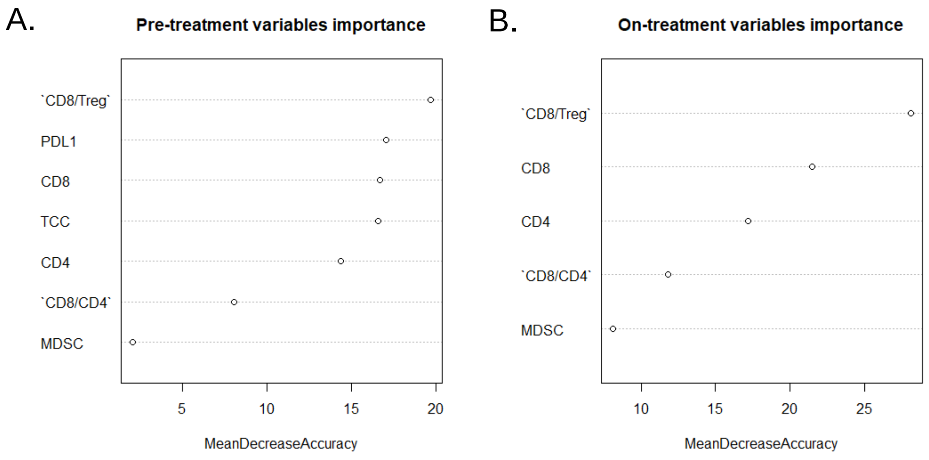


**Supplementary Figure 6. Variable importance calculated by random forest models of (A) pre-treatment and (B) on-treatment biomarkers of interest.** Treg, regulatory T cell. MDSC, myeloid-derived suppressor cells. TCC, the number of NSCLC-specific T cell clones.

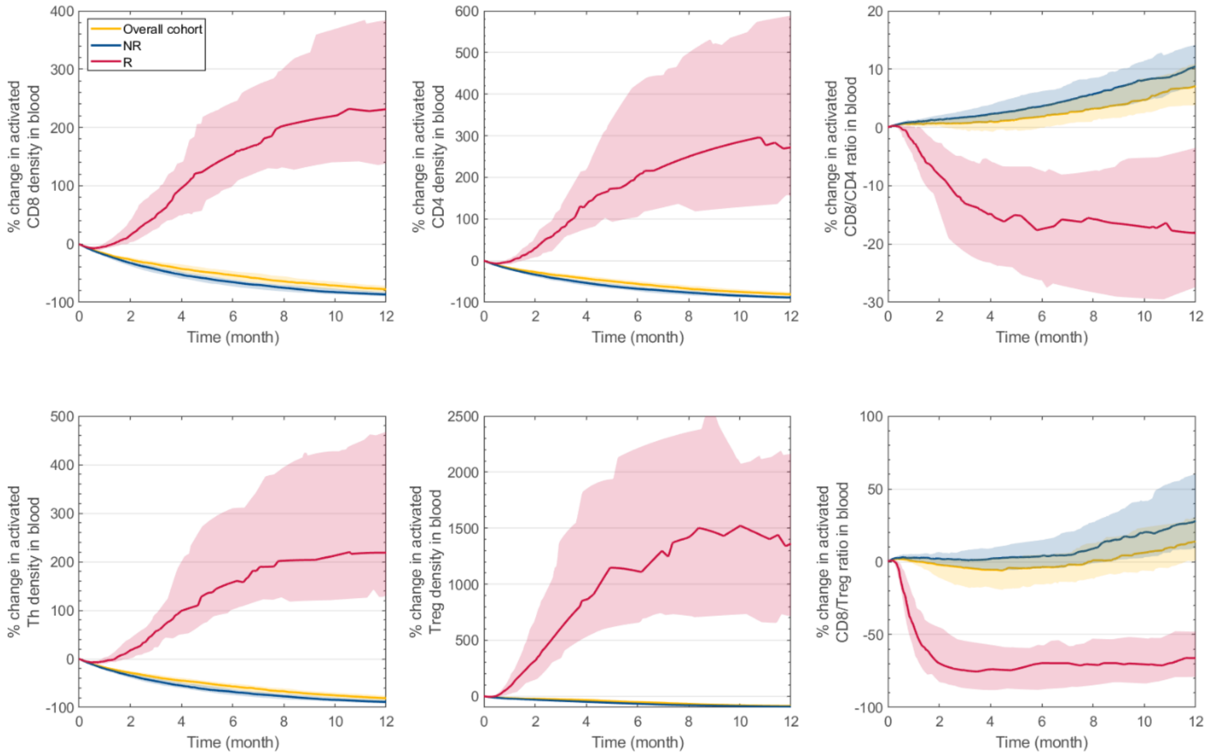


**Supplementary Figure 7. Effect of PD-L1 inhibition on model variables in the central compartment in responders, non-responders, and the overall virtual patient cohort, including percentage change in activated CD8 T cell, activated CD4 T cell, CD8/CD4 ratio, helper T cell (Th), regulatory T cell (Treg), and CD8/Treg ratio.** Solid lines represent median values. Shaded areas represent 95 percentile bootstrap confidence intervals.

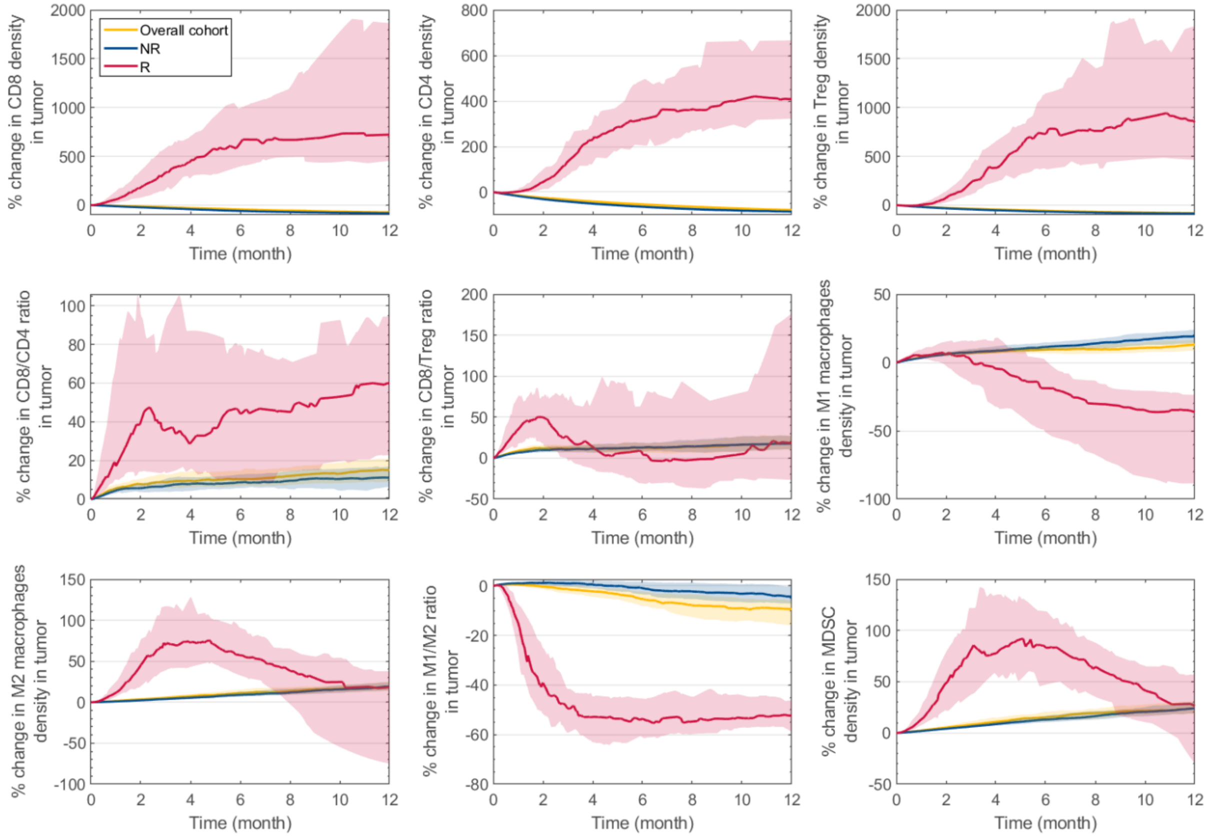


**Supplementary Figure 8. Effect of PD-L1 inhibition on model variables in the tumor compartment in responders (R), non-responders (NR), and all the virtual patients: percentage change in CD8 T cell, CD4 T cell, regulatory T cell (Treg), CD8/CD4 ratio, CD8/Treg ratio, M1 macrophages, M2 macrophages, M1/M2 ratio, and myeloid-derived suppressor cell (MDSC).** Solid lines represent median values. Shaded areas represent 95 percentile bootstrap confidence intervals.

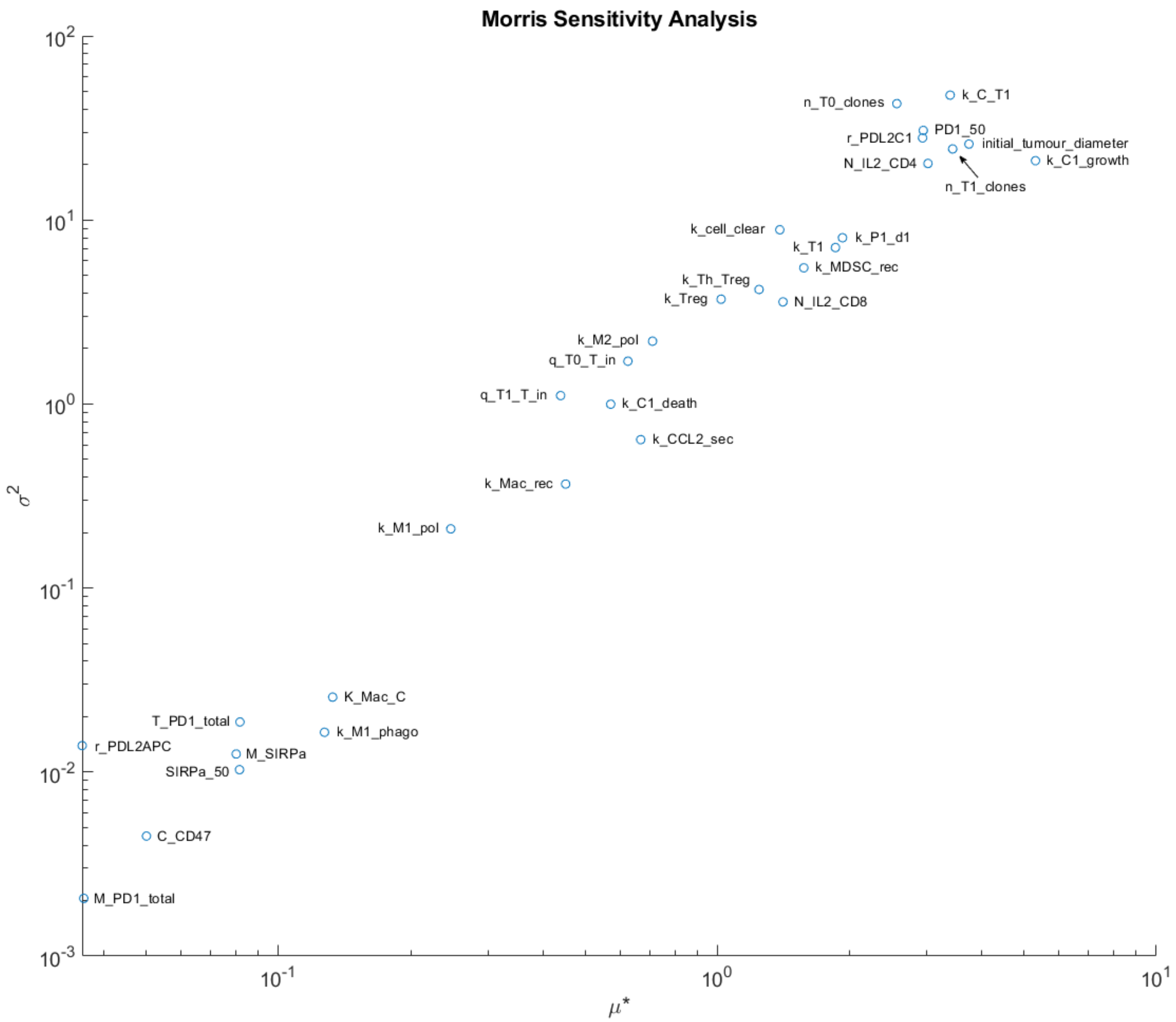


**Supplementary Figure 9. Global sensitivity analysis using Morris screening method.** Ranking of parameters determining tumor size at the end of durvalumab treatment (day 400). $\mu^*$, estimate of mean absolute value of elementary effects; $\sigma^2$, estimate of variance of elementary effects.
